# Correlation between the epigenetic modification of histone H3K9 acetylation of NR2B gene promoter in rat hippocampus and ethanol withdrawal syndrome

**DOI:** 10.1101/447771

**Authors:** Duan Li, Yanqing Zhang, Yanting Zhang, Qi Wang, Qin Miao, Yahui Xu, Jair C. Soares, Xiangyang Zhang, Ruiling Zhang

**Author notes:** Correspondence and requests for materials should be addressed to Xiang yang Zhang, Ruiling Zhang, Ph.D. The Second Affiliated Hospital, Xinxiang Medical University, P.R. China, Tel/Fax: ±86-0373-3373798;.

## Abstract

Our results showed that, in rat hippocampus, the ethanol withdrawal syndrome score was increased at 2 h, peaked at 6 h after withdrawal of ethanol, and reduced to the level parallel to the normal control group at day 3 after ethanol withdrawal. The NR2B mRNA expression and protein levels of rat hippocampus region showed similar patterns. Further correlation analyses indicted that both histone H3K9 acetylation in NR2B gene promoter and the expression levels of NR2B were positively associated with ethanol withdrawal syndrome. Chronic ethanol exposure may result in epigenetic modification of histone H3K9 acetylation in NR2B gene promoter in rat hippocampus, and the expression levels of NR2B were found to be positively correlated with EWS.

**Abstract:** Previous studies showed that an epigenetic modification of N-methyl-D-aspartate (NMDA) receptor, especially NMDA receptor 2B subunit (NR2B), was involved in the pathological process of ethanol withdrawal syndrome (EWS). However, the relationship between the epigenetic regulation of the NR2B gene in the rat hippocampus region and EWS were inconsistent. A rat model of chronic ethanol exposure was established. EWS score and the behavioral changes were recorded at different points in time. The NR2B expression levels and the histone H3K9 acetylation level in the NR2B gene promoter region were measured using qRT-PCR, Western blot, immunofluorescence and chromatin immunoprecipitation, respectively. Finally, the relationships between the epigenetic modification of histone H3K9 acetylation of NR2B gene promoter and EWS were examined. Our results showed that the EWS score was increased at 2 h, peaked at 6 h after withdrawal of ethanol, and reduced to the level parallel to the normal control group at day 3 after ethanol withdrawal. The NR2B mRNA expression and protein levels showed similar patterns. Further correlation analyses indicted that both histone H3K9 acetylation in NR2B gene promoter and the expression levels of NR2B were positively associated with EWS. Chronic ethanol exposure may result in epigenetic modification of histone H3K9 acetylation in NR2B gene promoter in rat hippocampus, and the expression levels of NR2B were found to be positively correlated with EWS.

## Introduction

Alcohol abuse can lead to the development of dependence and addiction, which may cause detrimental effects in brain development (Crews et al., 2007). The discontinuation of chronic ethanol exposure is associated with excitatory withdrawal signs called ethanol withdrawal syndrome (EWS). Patients withdrawing from ethanol present with autonomic hyperactivity, which may loss of appetite and seizures or evolve into hallucinations, anxiety, hyper excitability, convulsions and delirium tremens (Stehman and Mycyk, 2013). Although the signs of EWS in humans (Thompson, 1978) and rodents (Uzbay et al., 1997) have been well described, the mechanisms underlying physical dependence to ethanol and EWS are poorly understood.

Ethanol’s primary action on the central nervous system (CNS) is mediated by the inhibitory neurotransmitter γ-aminobutyric acid and the excitatory neurotransmitter glutamate, which binds to the N-methyl-D-aspartate (NMDA) receptor. NMDA receptors appear to be major cell-membrane bound targets for ethanol and may be responsible for mediating its long-term damaging effects (Dodd et al., 2004). Concurrently, ethanol also inhibits the excitatory action of glutamate at the NMDA receptor, leading to sedative and CNS depressing effects (Carlson et al., 2012). NMDA receptors, composed of NR1 combined with one or more NR2 (NR2A, NR2B, NR2C, NR2D) subunits and less commonly NR3 subunits (Kaniakova et al., 2012, Paoletti et al., 2013), are concentrated at postsynaptic sites (Liu et al., 1994). Increasing evidence is demonstrating the importance of the NR2B subunit in determining the pharmacological and functional properties of the NMDA receptor (Lett et al., 2014).

Chronic ethanol exposure leads to tolerance and physical and psychological dependence mediated by the up-regulation of the NMDA receptors(Carlson et al., 2012). NMDA receptors play a crucial role during chronic ethanol consumption and ethanol withdrawal and central reward systems (Self and Nestler, 1995). During chronic ethanol intoxication, the expression of NMDA receptors and their functions have been found to be up-regulated, possibly due to the depressing and antagonistic effects of ethanol on the NMDA receptors (Maler et al., 2005). These mechanisms may initiate neuroadaptive processes, causing the excitatory syndrome upon ethanol withdrawal (Maler et al., 2005).

In the brain, hippocampus plays a critical role in learning and memory and is particularly sensitive to ethanol exposure (Ryabinin, 1998). The region in the hippocampus expresses high levels of NMDA receptors (Goebel and Poosch, 1999, Monyer et al., 1994), which are severely impacted by ethanol abuse (Fadda and Rossetti, 1998, Jacobson et al., 1990). Hence, NMDA receptors at the site of hippocampus region are good candidates for studying excessive ethanol consumption and EWS. A previous study revealed increased numbers of NR2B subunits in the rat hippocampus region after 2 weeks of ethanol vapor exposure (Pian et al., 2010). There are epigenetic mechanisms involved in the development of addictive behavior (Warnault et al., 2013). Even though several reports have indicated that histone modifications are involved in alcohol-related events (Pandey et al., 2008), it is still unclear how this happens. It has been reported that ethanol exposure produces epigenetic modifications, including histone acetylation and methylation in hippocampus, striatum, and prefrontal cortex in both adult and adolescent rats (Moonat et al., 2013, Warnault et al., 2013). Other studies showed that histone acetylation has been examined widely, and it has been linked to the activation of gene transcription (Peterson and Laniel, 2004). These epigenetic modifications were also shown to regulate NR2B transcription in primary neuronal cultures (Qiang et al., 2010, Qiang et al., 2011). However, the functional role of the histone H3K9 acetylation in NR2B promoter region involved in EWS has yet to be established.

The purposes of the studies were, therefore, to examine the role of the epigenetic modification of NR2B gene in EWS, and the relationship between the expression of NR2B protein level and EWS in rat hippocampus region.

## Results

### Rat model for ethanol exposure

The average amount of 6% ethanol solution consumed everyday was 155.93 ml/kg body weight, which was equivalent to the intake of pure ethanol 7.39 g/kg body weight. There were no significant differences in ethanol consumption (g/Kg body weight) at every week and weeks of drinking (P>0.05) (Fig. S1A). After 16-week treatment, there were no significant differences in body weight between 5 ethanol consumption groups and normal control group (P>0.05), suggesting that low concentration of ethanol (6% in our current study) had no significant influence on body weight (Fig. S1B).

### Evaluation of ethanol withdrawal syndrome in rats

During the ethanol drinking period, all rats were more docile and less aggressive. After the ethanol was withdrawn, the rats showed the increased grooming, sneezing, irritability, tail stiffness, bowing of the back, and auditory epileptic seizures.

Based on different withdrawal time points, the EWS scores were 10.42±2.50 for 2 h, 15.42±1.93 for 6 h, 9.25±2.01 for 12 h, 7.67±1.92 for day 1, and 2.25±0.87 for day 3, respectively (Table 1). All these EWS scores were significantly higher than that of normal control group (1.50±0.80; all *P*<0.05). At 6 h time point, the EWS score was highest, and then decreased gradually to the level parallel to NC group (Fig. 1).

**Table 1.** Scores of ethanol withdrawal syndrome at each time point.

| Test of groups | stereotyped<br>behavior | irritability | tail stiffness | postural<br>malalignments | audiogenic<br>seizure | total scores |
| --- | --- | --- | --- | --- | --- | --- |
| *NC | 1.33 | 0.17 | 0 | 0 | 0 | 1.5 |
| **EW for 2 h | 3.5 | 0.42 | 3.17 | 2.25 | 1.08 | 10.4 |
| **EW for 6 h | 4.08 | 3.08 | 3.25 | 3.25 | 1.75 | 15.4 |
| **EW for 12 h | 2.67 | 1.33 | 2.33 | 2.25 | 0.67 | 9.25 |
| **EW for day 1 | 2.5 | 0.5 | 2 | 1.97 | 0.75 | 7.67 |
| **EW for day 3 | 1.5 | 0 | 0.5 | 0.08 | 0.17 | 2.25 |

Total rats are 48 rats, 8 rats per group. \*NC: Normal Control group; \*\*EW: Ethanol withdrawal group
Based on the previous report and a revised evaluation standard in Material and Methods, the EWS scores were counted including stereotyped behavior, irritability tail, stiffness, postural malalignments and audiogenic seizure at 0 h, 2 h, 6 h, 1 h, day 1 and day 3 after ethanol withdrawal, respectively.

**Fig.1.**
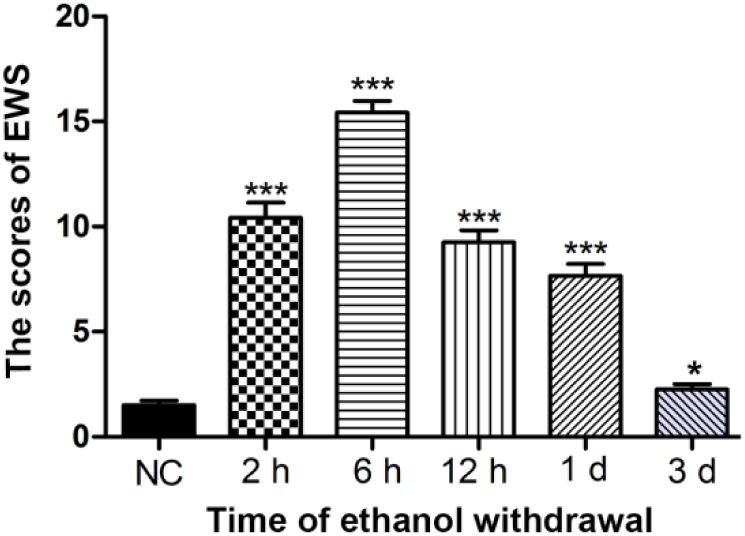
Evaluation of ethanol withdrawal syndrome in rats after removal of ethanol. The ethanol withdrawal syndrome (EWS) scores were counted at the different time points after ethanol withdrawal. Data are expressed as mean ± SEM (n = 8 rats per group). *indicates the comparisons to the normal control (NC) group; \**P*<0.05, *** *P*<0.001.

### Expression level of NR2B protein in hippocampus at different withdrawal time points

The total protein and total RNA from rat hippocampus were extracted separately. The relative expression of the NR2B mRNA was 0.33 ± 0.046 for 2 h, 0.242±0.059 for 6 h, 0.202±0.044 for 12 h, 0.179±0.039 for day 1 and 0.154 ± 0.022 for day 3, which were all significantly higher than that in normal control group (0.13±0.022; all *P*<0.05). The expression of NR2B mRNA peaked at 2 h after withdrawal and the amount of expression began to decrease at the subsequent time points (Fig. 2A).

**Fig.2.**
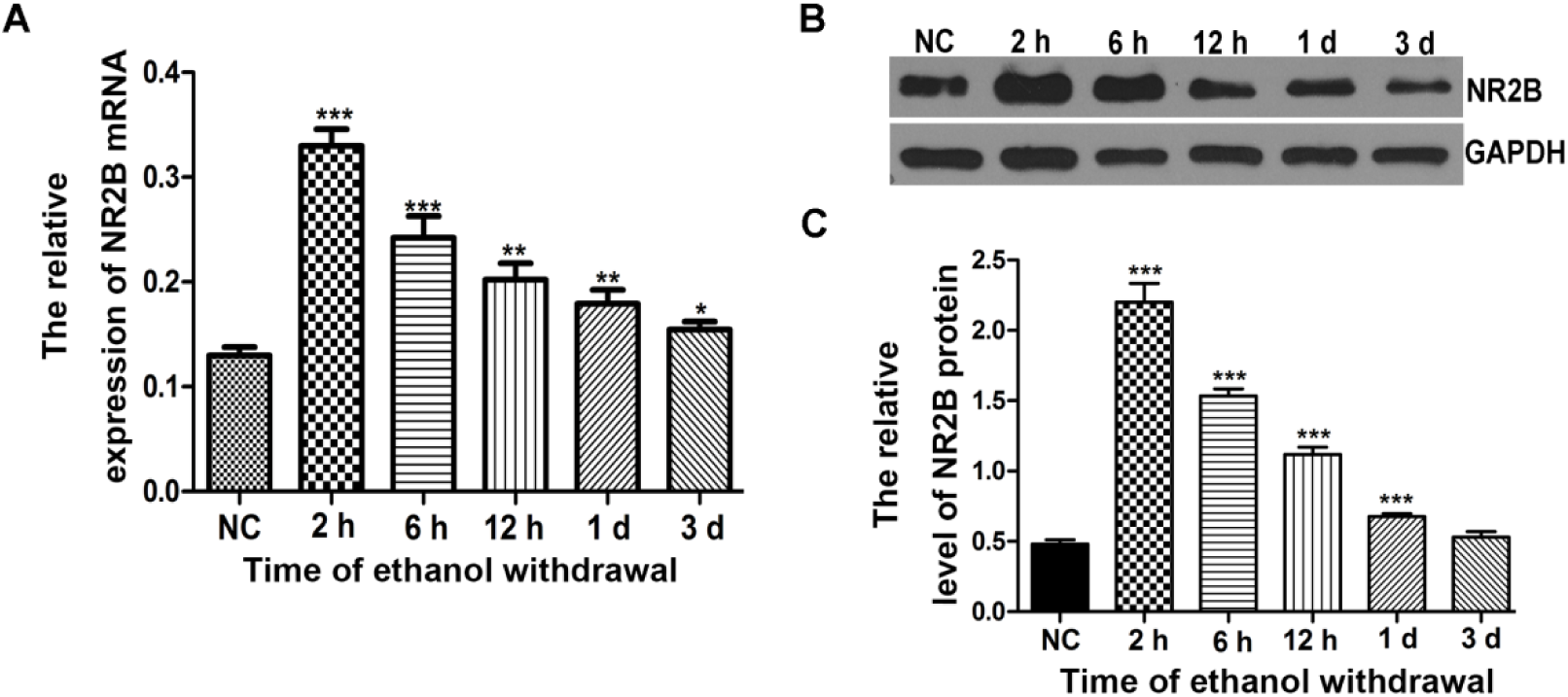
The expression levels of NR2B gene in rat hippocampus at different time points after ethanol withdrawal. **A**. Relative mRNA of NR2B expression at 2 h, 6 h, 12 h, 1 d and 3 d after ethanol withdrawal were assayed through qRT-PCR, compared with normal control group. Data are expressed as mean ± SEM (n = 8 rats per group). **B**. The expression levels of NR2B protein in rat hippocampus region were assayed by the Western blot at different time points after ethanol withdrawal. Data are expressed as mean ± SEM (n = 8 rats per group). **C**. The relative expression levels of NR2B protein were presented in the form of a histogram at different time points after ethanol withdrawal by gray scale analysis. *indicates the comparisons to the normal control (NC) group; *P>0.05, *** *P*<0.001.

With Western blot assay, the level of NR2B protein was examined at different withdrawal time point. The ratio of NR2B gray level to GAPDH gray level represented the relative level of NR2B protein, and this level were 2.201±0.375 for 2 h, 1.535±0.137 for 6 h, 1.117±0.149 for 12 h, 0.677±0.054 for day 1 and 0.53±0.113 for day 3, which were all significantly higher than that in normal control group (0.481±0.085; all *P*<0.05), except for that at day 3 (*P* >0.05) (Fig. 2B). The protein level of NR2B peaked at 2 h after withdrawal and had decreased to normal level at day 3 (Fig. 2C).

### Immunofluorescence assay for the NR2B protein in rat hippocampus

The level of NR2B protein at the tissue level was assaying using immunofluorescence assay. Figure 3A showed that the protein level of NR2B peaked at 2 h after withdrawal and had decreased to the level of normal control group at day 3. The NR2B protein level by the immunofluorescence intensity displayed the same pattern as that by Western blot assay, with 1411.36±210.72 for 2 h, 987.38±104.32 for 6 h, 823.62±48.14 for 12 h, 554.71±73.13 for day 1 and 437.24±123.76 for day 3, which were all significantly higher than that in the normal control group (378.56±67.65; all *P*<0.05), except for that at day 3 (P>0.05) (Fig.3B).

**Fig.3.**
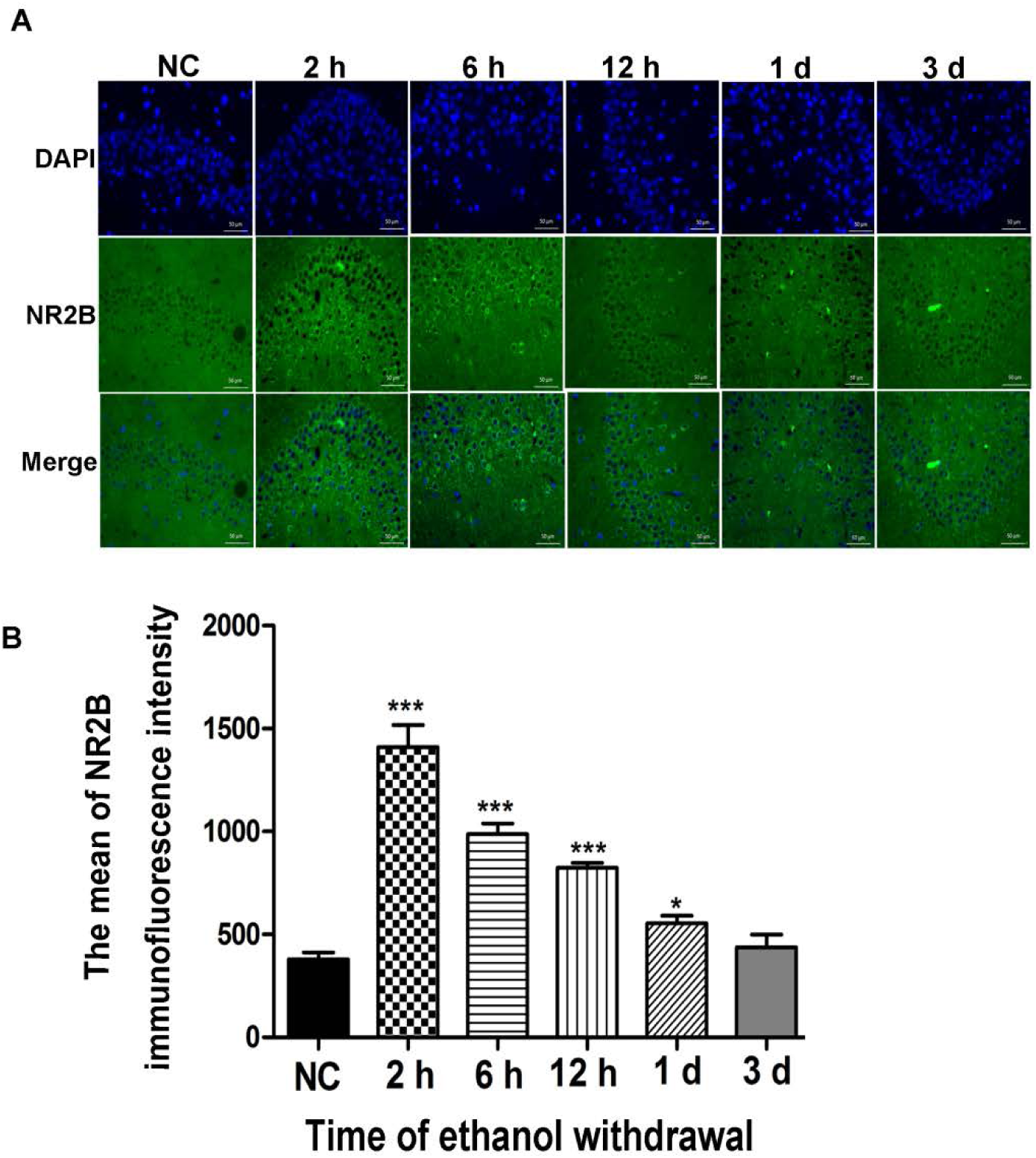
The expression of NR2B protein in rat hippocampus at different withdrawal time points by immunofluorescence assay. **A**. NR2B expression in hippocampus region at 2 h, 6 h, 12 h, 1 d and 3 d after ethanol withdrawal was assessed by the immunofluorescence assay. The nuclei were indicated by DAPI staining (blue) and NR2B protein was stained by fluorescent antibody (green). Scale bars: 30μm. Data are expressed as mean ± SEM. **B**. NR2B immunofluorescence intensity at 2 h, 6 h, 12 h, 1 d and 3 d after ethanol withdrawal were represented by histogram, compared with normal control group. *indicates the comparisons to the normal control (NC) group; \**P*<0.05, *** *P*<0.001.

### H3K9 levels of NR2B gene in hippocampus at different withdrawal time points

The levels of histone H3K9 acetylation of NR2B promoter region in the hippocampus were measured by CHIP. According to the experimental instructions, the fold enrichment of relative acetylation represents the percent of the H3K9 acetylation of NR2B gene at the time point of ethanol withdrawal to at the input. Figure 4 showed that the fold enrichment of histone H3K9 acetylation of NR2B gene to input was 133.38±6.947, 121.85±3.643, 123.0±6.053, 122.21±5.661 and 93.20±3.133 after the ethanol withdrawal at 2 h, 6 h, 12 h, day 1 and day 3, respectively, while the fold enrichment to input was 123.19±5.107 for the normal control group. In this way the histone H3K9 acetylation of NR2B gene at 2 h after ethanol withdrawal was significantly higher than in the normal control group (*P*<0.001), while the fold enrichment of NR2B gene acetylation gradually decreased, showing significant differences at others time point (all P<0.05) except for day 3 (Fig. 4).

**Fig.4.**
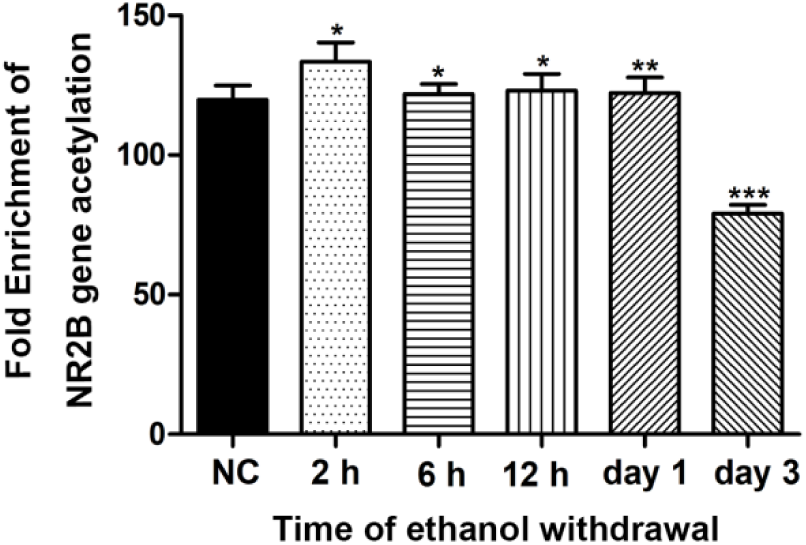
H3K9 levels of NR2B gene in rat hippocampus at different time points after ethanol withdrawal using Chromatin immunoprecipitation. The level of histone H3K9 acetylation of NR2B gene promoter in rat hippocampus region at the different time points of 2 h, 6 h, 12 h, 1 d and 3 d after ethanol withdrawal was assessed by the Chromatin immunoprecipitation. Data are expressed as mean ± SEM (n = 8 rats per group). *indicates the comparsions to the normal control (NC) group; * *P* <0.05, *** *P* <0.001.

### Correlation between the EWS score, the levels of H3K9 acetylation of NR2B gene promoter and the expression of NR2B in rat hippocampus

Correlation analysis showed the expression levels of the hippocampus NR2B protein to be positively correlated with the EWS scores (*r*=0.522, df=40, *P*=0.0006) (Fig. 5A). Similarly, there was a significantly positive correlation between the expression of NR2B mRNA and the level of NR2B protein (*r*=0.7754, df=40, *P*<0.0001) (Fig. 5B). These results suggested a close relationship between EWS and the expression of NR2B mRNA and protein.

**Fig.5.**
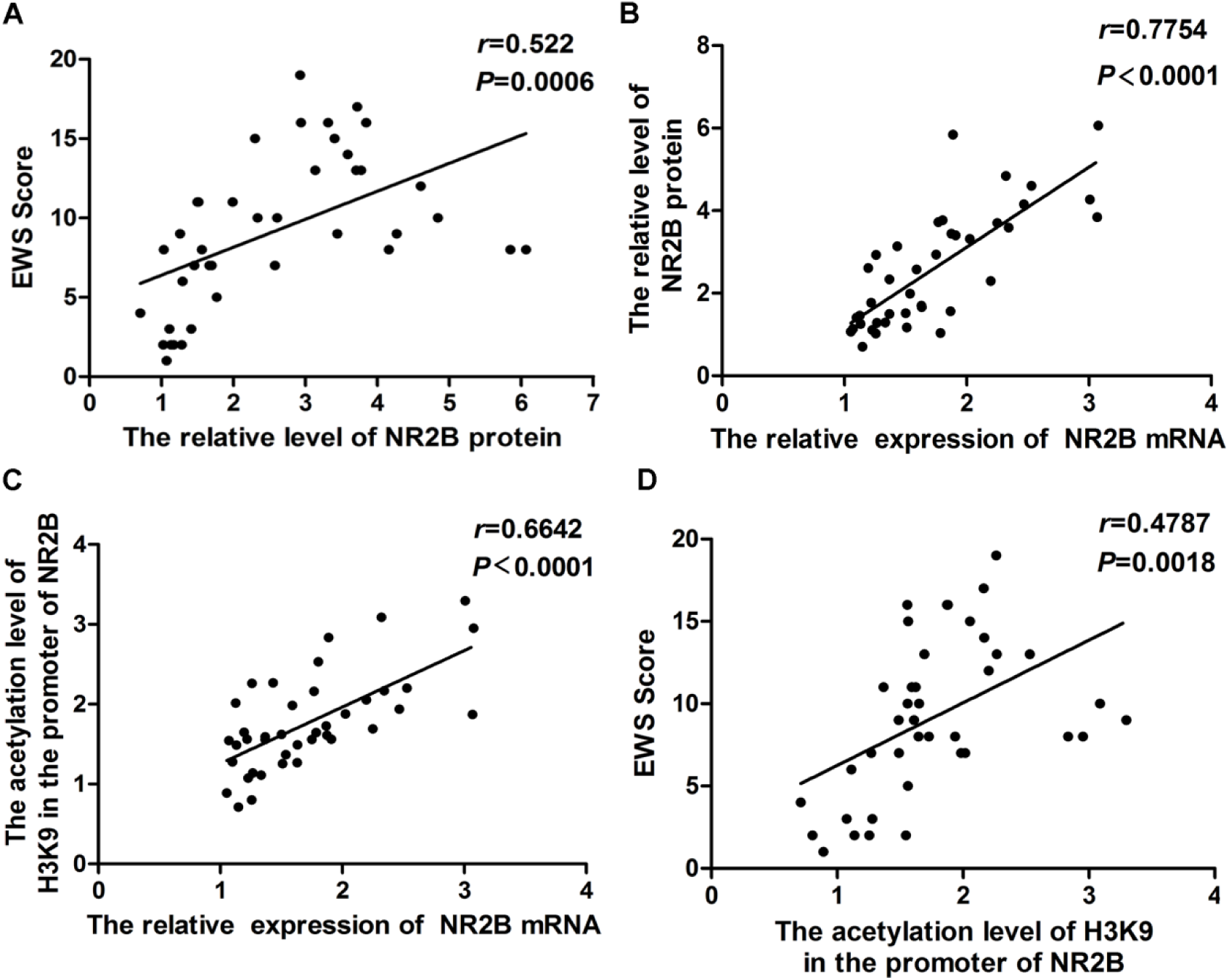
The correlation between the ethanol withdrawal syndrome score, the levels of H3K9 acetylation of NR2B gene promoter and the expression of NR2B in rat hippocampus. **A**. There was a significantly positive correlation between EWS score and the relative level of NR2B protein (*r*=0.522, *P* =0.0006). **B**. There was a significantly positive correlation between the relative level of NR2B protein and the relative expression of NR2B mRNA (*r*=0.7754, *P* <0.0001). **C**. There was a significantly positive correlation between the acetylation level of H3K9 in the promoter of NR2B and the relative expression of NR2B mRNA (*r*=0.6642, *P* <0.0001). **D**. There was a significantly positive correlation between EWS score and the histone H3K9 acetylation of in the promoter of NR2B gene (*r*=0.4787, *P* =0.0018). Note: The pooled data from the ethanol withdrawal groups were used for this analysis (n=40, 8 rats per group) in comparison with normal control group.

Subsequent correlation analysis showed that the levels of histone H3K9 acetylation in NR2B promoter region to be positively correlated with both the expression of NR2B mRNA (*r*=0.6642, df=40, *P*<0.0001 (Fig. 5C) and the EWS scores (*r*=0.4787, df=40, *P*=0.0018 (Fig. 5D).

## Discussion

EWS consists of a variety of behavioral symptoms, physiological signs and the electrophysiological changes in the central nervous system (Thompson, 1978). NMDA receptor is an ionotropic receptor that is involved in synaptic plasticity and memory formation, which is the main target of ethanol in the CNS (Chandrasekar, 2013, Whitlock et al., 2006). Although the effects of ethanol on the NMDA receptor have still been unclear, it has been found that the effects of ethanol on CNS are related to the NMDA receptor subunits, especially NR2B (Brady et al., 2013). Previous studies showed that many neurological symptoms caused by ethanol, such as ethanol withdrawal symptoms and nerve excitotoxicity, are closely related to NR2B subunit (Brady et al., 2013). For example, a previous study reported that the expression level of NR2B protein was significantly higher in cultured hippocampus and cortical primary neurons of rats after chronic ethanol treatment, while NR2A, NR2C and NR2D did not change significantly (Pian et al., 2010). Also, in the animal experiment performed in rats, the level of NR2B mRNA was significantly higher after 2 weeks of chronic ethanol treatment (Roberto et al., 2006). Our results indicated that NR2B protein was higher after ethanol withdrawal for 2 h. Taken together, these findings suggest that the expression level of NR2B protein in hippocampus may be involved in ethanol abuse/dependence and withdrawal behaviors.

In hippocampal region of adult male Wistar rats, NR2B was significantly increased after chronic ethanol exposure (Law et al., 2003). Moreover, there was a notable correlation between the relative mRNA expression of NR2B and the level of NR2B protein. Here, we speculated that there was correlation between EWS and the relative expression of NR2B mRNA.

Histone acetylation includes acetylation of H3 and H4. H3K9 is the only residue that can be acetylated or methylated (Nakayama et al., 2001). Histone H3K9 acetylation was found to be inversely correlated with H3K9 methylation (Yamada et al., 2008). A previous study found that acute ethanol use induced the significantly increased histone H3K9 acetylation, with less change in K14, K18 and K23 acetylation status (Park et al., 2005). One recent study found that chronic intermittent ethanol exposure and ethanol withdrawal increased the expression of NR2B and the acetylation level of H3K9 in cultured mouse cortical neurons (Qiang et al., 2011). These results suggest that ethanol may selectively produce the effects on the histone H3K9 as the target.

There are several limitations in this study. First, the role of epigenetic modification of NR2B gene promoter in ethanol withdrawal syndrome was examined solely in hippocampus. Whether such a relationship exists in other parts of the brain merits further investigation. Second, the underlying mechanisms that EWS caused in histone acetylation of NR2B is still unclear, so it will need to be examined in further research.

Taken together, it is reasonable to speculate that ethanol withdrawal may lead to the increased levels of histone H3K9 acetylation of NR2B gene and the NR2B protein expression in hippocampus region, which may play an important role in the development of EWS.

## Materials and Methods

### Preparation for the rat model of ethanol exposure

The animal experiment was performed in the Key Psychiatric Laboratory of the Second Affiliated Hospital of Xinxiang Medical University between April 2016 and February 2017. Care of all animals was carried out in accordance with the Institutional Animal Care and Use Committee of Xinxiang Medical University. All experiments included in the present report were replicated using at least two different rat litters.

Forty-eight male Wistar rats weighing 180-220g were provided by the Beijing Vital River Laboratory Animal Technology Co., Ltd., China (license No: SCXK (Jing) 2012-0001). All rats were group-housed with free access to water and food in the vivarium, which had a 12-hour light/dark cycle and a temperature regulated environment. The 40 rats in ethanol withdrawal groups were administered with drinking the concentration of 6% ethanol-containing liquid (v/v) for 16 weeks, while rats in normal control group (NC, 8 rats) were maintained with water for 16 weeks.

### Scores of ethanol withdrawal syndrome (EWS)

At the end of 6% ethanol-containing liquid diet administration, ethanol was withdrawal from the diet by replacing the diet without ethanol. The 40 ethanol-exposure rats were assigned into 5 different time point groups at random (n=8 for each group). According to the previous report (Erden et al., 1999) and a revised evaluation standard of EWS (Supplements Table1), the EWS was tested and scored at 2 h, 6 h, 1 h, day 1 and day 3 after ethanol withdrawal, respectively.

### Preparation for tissue samples

At the end of each withdrawal time point, all of the animals were killed after anesthesia with an injection of chloral hydrate after the behavioral features of EWS were recorded. The hippocampus was removed from the brain and quickly frozen in liquid nitrogen for further experiments.

### Western blot for NR2B protein and RT-PCR for NR2B mRNA

Hippocampus tissues were extracted and total proteins were used for Western blot as previously described (Liang et al., 2014). Total RNA of the hippocampus tissues was extracted by Trizol Reagent (Invitrogen) according to the manufacturer’s instructions. The RT-PCR for NR2B mRNA were performed with a QuantiFast SYBR green RT-PCR kit (Qiagen, DüsseLDsorf, Germany) and reaction conditions were carried out according to the manufacturer’s instructions (Applied Biosystems, Foster City, CA). GAPDH expression was an endogenous control to normalize the data. The primer sequence of NR2B gene and GAPDH served as endogenous control. They were listed in Supplements Table 2. All of the quantifications were performed with an ABI Prism 7500 system (Applied Biosystems).

### Immunofluorescence assay for NR2B protein of rat hippocampus

For NR2B immunofluorescence, paraffin sections were placed in an oven at 60°C for 30 min. The sections were then deparaffinized and rehydrated by xylene twice and gradient ethanol from pure to 70%. Heat-induced retrieval of antigen was performed using 10mM sodium citrate (pH 6.0), 3% hydrogen peroxide for 10 min to suppress the endogenous peroxidase activity, and 1% bovine serum albumin (all from Sigma, St. Louis, MO, US) for 15 min to block nonspecific binding. After this, the sections were incubated in rabbit polyclonal NR2B antibody overnight at 4°C. After washing three times in PBS, 10 min each time, the sections were incubated with rabbit FITC-labeled fluorescent antibody (1:400) without light at room temperature for 2 h. Nuclei were stained with 4’, 6-diamidino-2-phenylindole (DAPI) without light at room temperature for 10 min. Four random fields in each group were observed under a Leica TCS-SL confocal microscope. The fluorescence intensity was quantified with Image J software (NIH).

### Chromatin immunoprecipitation (ChIP) for hippocampal NR2B gene promoter

The ChIP procedure was performed using the instructions provided with the ChIP kit (Millipore, Bedford, MA, US) and with anti-acetylated histone H3 at Lys9 (ac-H3K9) antibody (Rabitt polyclonal, Abcam,) or IgG (Santa Cruz Biotechnology) as negative control. We designed the special primers of NR2B gene promoter region to amplify, GAPDH as the internal control (sequences listed in Supplements Table 3) and the immunoprecipitated DNA, input DNA and negative control DNA served as templates for qRT-PCR. The levels of histone acetylation in the promoter area of the NR2B gene were determined by the fold enrichment. The fold enrichment was calculated relative to the percentage of input DNA.

### Statistical analysis

All of the data were shown as mean ± SEM and were analyzed utilizing SPSS 15.0 statistical software (SPSS, Chicago, IL, US). One-way analysis of variance (ANOVA) was used to make the comparisons between different ethanol withdrawal groups and normal control group. Pearson’s correlation test of the five groups of alcohol abstinence groups was used to assess the relationships between EWS scores, the expression level of NR2B protein, the mRNA expression of NR2B gene and the histone H3K9 acetylation of NR2B gene in hippocampus region. Statistical significance level was set to *P* < 0.05.

## Acknowledgements

Duan Li and Ruiling Zhang are responsible for the study concept and design. Yanqing Zhang, Yanting Zhang, Qi Wang, Qin Miao and Yahui Xu carried out all animal experiments and part of the molecular biology experiment. Duan Li and Ruiling Zhang carried on the molecular biology experiment and the data statistical analysis. Xiangyang Zhang and Jair C.Soares carried out the language polish of the article. Duan Li and Ruiling Zhang were in charge of data analysis and interpretation and manuscript drafting. All authors critically reviewed content and approved final version for publication.

## Compelting of interest

The authors declare that they have no conflict of interest.

## Abbreviations

EWS: ethanol withdrawal syndrome
NMDA: N-methyl-D-aspartate
NR2B: N-methyl-D-aspartate receptor 2 B
ChIP: Chromatin immunoprecipitation
qRT-PCR: quantitative Real time PCR
CIE: chronic intermittent ethanol

